# Experimental evidence of nocturnal pollination services for berry crops

**DOI:** 10.1101/2023.11.02.565389

**Authors:** Elsa Blareau, Fabrice Requier

**Author notes:** Correspondence: F. Requier.

## Abstract

Insect-mediated pollination is essential for crop production but is mainly studied considering diurnal pollinators only. Using pollinator exclusion techniques to prevent either diurnal or nocturnal insect visits we estimate the contribution of day-active and night-active pollinators to fruit set, seed set, fruit weight and fruit volume in three berry crops. We find that nocturnal pollination improves seed set compared to exclusion of all insect visits for one crop (raspberry), confirming the existence and importance of a nocturnal pollination service. Moreover, we found a general pattern where best crop pollination resulted from combined services of diurnal and nocturnal insect visits. These results call for urgent consideration of nocturnal pollination in crop production and management, towards inclusive pollinator-friendly schemes including night-active pollinators.

## Introduction

Animal pollination, the processes by which an animal facilitates the transfer of pollen from the anthers of one flower to the stigma of another, is essential for the reproduction of 87.5% of all flowering plants [1]. Although some crops, such as cereals, only rely on wind for pollination, animals are involved in the pollination of over half of the leading crops worldwide [2]. The crop pollination service is valued at 195-387 billion US dollars annually [3]. In agriculture, animal pollination has implications for food security, since many animal pollinated crops contain nutrients required for a healthy human diet [4–6]. In addition, pollination has value in areas such as medicine, the production of biofuels or even the reproduction of aesthetically important plants [4,7].

Among animal pollinators, insects play a critical role in agriculture. Insect pollinators come from the orders Hymenoptera (bees and wasps), Diptera (flies), in particular flies of the family Syrphidae (hoverflies), Coleoptera (beetles) and Lepidoptera (butterflies and moths) [8,9]. The Western honey bee *Apis mellifera* is known as an economically important pollinator in agriculture, since this species is managed by beekeepers, who place beehives in close proximity to crops to make sure the pollination service is provided [10]. However, wild insect pollinators provide significant and essential contributions to crop pollination [8,11], as do some non-insect pollinators in the tropics, such as birds and bats [12,13].

The evidence for a decline of insect pollinators [14–16] raises alarm for the resilience of their supporting pollination services to pollinator-dependent crops [7]. Indeed, less diverse biological communities are often thought to provide ecosystem services that are less resilient to disturbance over space or time [17,18]. Crop pollination is therefore commonly cited as an example of an endangered ecosystem service, with evidence of pollination deficits (e.g. for apple [19,20] and for highbush blueberry [21]), particularly in intensive agricultural landscapes [22] and their relations to global environmental changes that affect pollinator biodiversity [15]. Pollinator-friendly schemes are thus recommended to protect bee populations in agricultural landscapes practices [10,23] such as restoring semi-natural habitats [24], planting flowers [25], providing nesting habitats [26] and reducing intensive farming practices [27].

However, most of our knowledge about pollination and pollinators focus on diurnal activity, neglecting the contribution of nocturnal pollinators to crop production [13,28]. Night-active pollinators include vertebrates such as bats (under tropical climates) as well as invertebrates such as moths, beetles and some bee species [28–36]. Some studies have investigated whether nocturnal pollinators contribute to crop pollination. Firstly, nocturnal pollinators were observed visiting flowers of numerous tropical and temperate crops [37]. Interestingly, recent studies showed nocturnal pollinators facilitating pollen transfer in avocado crops [28] and contributing to fruit set in apple orchards [38] and strawberry crops [39].

In this study, we explore whether nocturnal pollination services can be observed in other crops, to discuss their inclusion in current pollinator-friendly schemes. To achieve this, we studied fruit set, seed set, fruit weight and fruit volume for different pollination scenarios in three berry crops: raspberry (*Rubus idaeus*), red currant (*Ribes rubrum*) and black currant (*Ribes nigrum*). These crops are known to benefit from animal pollination [2], but the contribution of nocturnal pollination is yet to be assessed. We designed experiments using pollinator exclusion techniques to prevent either diurnal or nocturnal insect visits in order to assess the potential contribution of day-active and night-active pollinators.

## Materials and methods

### Experimental design

The experiment was carried out in the protected area *Parc Naturel Régional Périgord-Limousin* in western France (45°34’16.2"N 0°55’05.5"E), on privately owned land, in 2020 (from March to July). The land cover was composed of a mixed landscape area with 37.71% of agricultural land and 38.12% of forested area in a 1.5km radius [40]. Having a study site in a mixed habitat, in a protected area, allowed for us to examine the contribution of nocturnal pollinators in favourable conditions for pollinator diversity, as opposed to intensive agricultural landscapes, where unfavourable conditions potentially restricting the presence of nocturnal pollinators may prevent us from detecting whether they contribute to pollination services. The three study crops were raspberry (*Rubus idaeus,* var. Zeva), red currant (*Ribes rubrum, var.* Versaillaise Rouge) and black currant (*Ribes nigrum, var.* Black Down). Yields are known to decrease in the absence of pollinators for raspberry [41,42], with higher pollinator dependency for raspberry than red currant [2]. In black currant, flowers can be qualified as self-compatible [43], but animal pollination increases yield of this crop [44].

### Diurnal pollinator observations

Diurnal pollinator observation sessions were performed between the 3rd of March 2020 and the 30th of April 2020. Raspberry and black currant flowers were observed at 17 occasions, and red currant flowers were observed 18 times. A total of 147 raspberry flowers (average of 8.65 flowers per session) were observed, 552 red currant flowers (average of 30.67 flowers per session) and 756 black currant flowers (average of 44.47 flowers per session). Each observation session lasted 5 minutes. Pollinators observed were sorted into the following nine categories: honey bee (Apis mellifera), bumble bee (Bombus sp.), wild bee, hoverfly (Syrphidae), other fly (Diptera), wasp (Vespidae), ant (Formicidae), beetle (Coleoptera) and spider (Aranea).

### Pollination treatments

In march 2020, flowers from two plants of each of these three crop species were affected to different pollination treatments. This design was chosen since only two plants of each crop were available for the study. All treatments were performed on both plants. For raspberry, 54 flowers were used (14±3 per treatment), for red currant 134 flowers were considered (34±14 per treatment) and for black currant there were 141 flowers (35±6 flowers per treatment). For red currant and black currant, flowers develop in racemes. Flowers within the same raceme were allocated to the same treatment and considered independently. In addition, several racemes per plant were allocated to each treatment. The four pollination treatments were: flowers excluded from pollinator visits (Closed), flowers open to pollinator visits only during the day (Day) and only during the night (Night), and flowers open to pollinator visits during both day and night (Day+Night). Pollinator exclusion was achieved by bagging flowers with mesh netting, preventing any insect visits (Alt’Droso Maraichage, 0.8 × 0.8 mm mesh). Flowers allocated to the Day treatment were bagged at sunrise (7:30 am) and exposed to pollinators for 11 hours and 30 minutes per day. Those allocated to the Night treatment were bagged at sunset (7:00 pm) and were exposed to pollinators for 12 hours and 30 minutes per day. Flowers allocated to the Day+Night treatment were exposed continuously to pollinators. Comparing the Closed and Day+Night treatments measures pollination service provided by all pollinators [20], meaning the additive contributions of diurnal and nocturnal pollinators. Comparing the Closed and Day treatments measures the pollination service provided by diurnal pollinators only. On the other hand, comparing the Closed and Night treatments measures the pollination service provided by nocturnal pollinators only.

### Fruit yield measures

Once flowering was over and fruits began to form, we measured fruit set of all three crops on a binary scale, recording for each flower the success (1) or failure (0) of fruit production. Fruits from all three crops were harvested over June and July 2020. We measured fruit weight (Brifit, Digital Kitchen Scale, accuracy 0.01g, capacity 500g). We estimated fruit volume using two measures of fruit size: maximum height and width (Mitutoyo, Digimatic Caliper, accuracy 0.01mm, capacity 150mm). Assuming the shape of a raspberry is half of an ellipsoid, we used the following formula to calculate fruit volume:

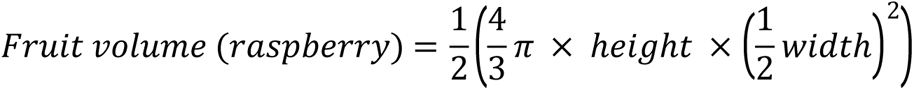

Red currant and black currant volumes were computed assuming fruits are shaped like spheres:

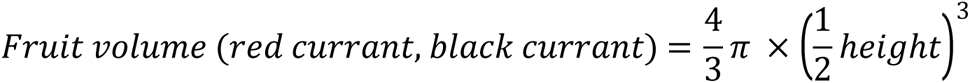

In addition, we measured seed set by counting the number of seeds produced by each fruit. These metrics are not linked to commercial standards [45] but were chosen since they are known to respond to pollination treatments for blueberry (fruit set, weight [21], and fruit size [46] and raspberry (fruit height and drupelet/seed set [47]).

### Statistical analysis

The statistical analysis was performed in R 4.2.2 [48]. The packages “stats” [48], dplyr [49] and “multcomp” [50] were used for the analysis. The package “ggplot2” [51] was used for graphical representation of data. We tested the effects of pollination treatments on fruit set using a generalised linear model (GLM) with binomial error structure, and a GLM with Poisson error structure for seed set. After checking for normality using a Shapiro-Wilk test, we used linear models (LM) to test the effects of pollination treatments on fruit weight and fruit volume. We used Tukey post-hoc tests for each model, to compare pairwise effects of pollination treatments. We considered a threshold of 5% to indicate significant differences between groups.

## Results

### Diurnal pollinator observations

We observed 109 diurnal visits on raspberry flowers in total (frequency of 0.39 visits per flower per minute), with ants as main diurnal visitors (67 visits), followed by bumble bees (25 visits), honey bees (15 visits), hoverflies (1 visit) and spiders (1 visit). We observed 7 visits to red currant flowers (frequency of 0.01 visits per flower per minute), all flies. We observed 23 visits to black currant flowers (frequency of 0.02 visits per flower per minute). These were predominantly flies (10 visits), followed by ants (6 visits), bumble bees (5 visits), wasps (4 visits), beetles (2 visits), wild bees (1 visit).

### Fruit set

We found a general pattern for all three crops where fruit set was minimal for the Closed pollination treatment, then increased successively in the Night, then the Day treatments, and reached a maximum for the Day+Night treatment (Fig 1). However, this general pattern differed among crops in the values of fruit set and in the intensity of the treatment effects. Raspberry showed the highest fruit set with 81.5% of flowers (n=44/54) giving a fruit, without significant differences between pollination treatments (Fig 1a; Supplementary information, S1 Table). Fruit set was intermediate in red currant with 47% of flowers producing a fruit (n=63/134). Fruit set for this crop was significantly higher in the Day treatment compared to the Closed treatment (p=0.005; Fig 1b, S1 Table). Fruit set was also higher in the Day+Night treatment compared to the Closed treatment (p<0.001; Fig 1b, S1 Table), and compared to the Night treatment (p=0.008; Fig 1b, S1 Table). For black currant, only 18.4% of flowers successfully produced a fruit (n=26/141), without significant differences in fruit set between pollination treatments (Fig 1c, S1 Table).

**Figure 1.**
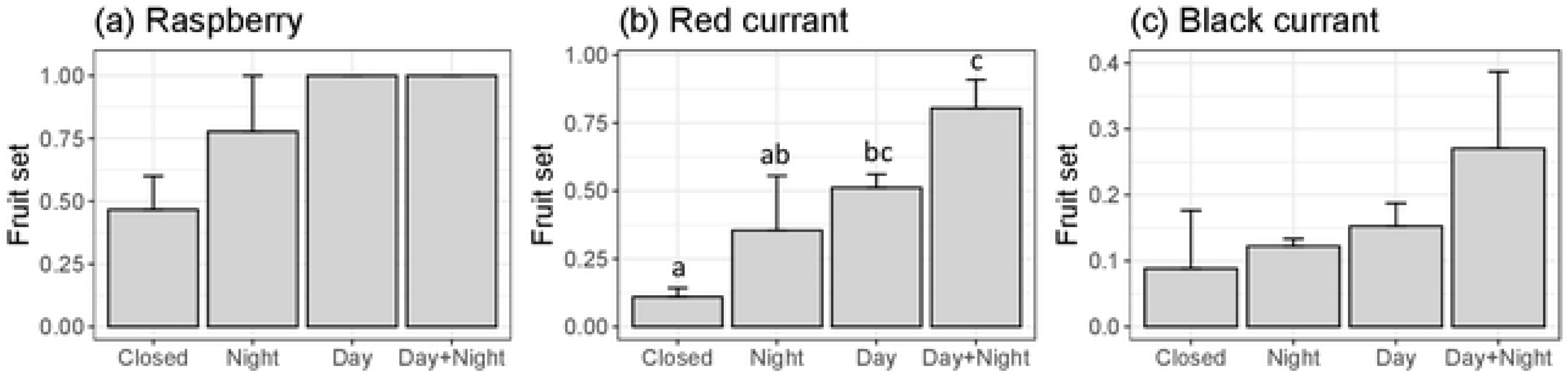
Comparing fruit set (mean+se) between Closed, Night, Day and Day+Night pollination treatments for (**a**) raspberry, (**b**) red currant, and (**c**) black currant. Different small letters indicate significant differences with a risk of 5%.

### Fruit quality

For flowers that successfully produced a fruit, we found a similar pattern of seed set as fruit set for raspberry and red currant, with lowest seed set for the Closed pollination treatment, then successive increases for the Night, Day and Day+Night pollination treatments. Once again, this pattern differs between these crops in seed set values and the intensity of pollination treatment effect. Seed set was higher in the Night treatment compared to the Closed treatment in raspberry (p<0.001; n=44 fruits; average of 49±23 seeds per fruit; Fig 2a, S2 Table). For raspberry, seed set was also higher in Day and Day+Night treatments compared to the Closed treatment (p<0.001; Fig 2a, S2 Table), and also higher in Day and Day+Night treatments compared to the Night treatment (p<0.001; Fig 2a, S2 Table). The average seed set for red currant was 7±5 per fruit (n=63 fruits). We find that seed set was higher in the Day treatment compared to the Closed treatment for this crop (p=0.016; Fig 2b, S2 Table), while the other pollination treatments did not differ. Average seed set for black currant was 8±5 per fruit (n=26 fruits). Conversely to the other crops, seed set was significantly higher in the Closed treatment compared to the Day+Night treatment (p<0.001; Fig 2c, S2 Table). Seed set was also higher in the Night treatment compared to the Day+Night treatment (p=0.021; Fig 2c, S2 Table). We did not find any significant difference of fruit weight and volume between pollination treatments for any of the three crops (S1 and S2 Figs, S3 and S4 Tables).

**Figure 2.**
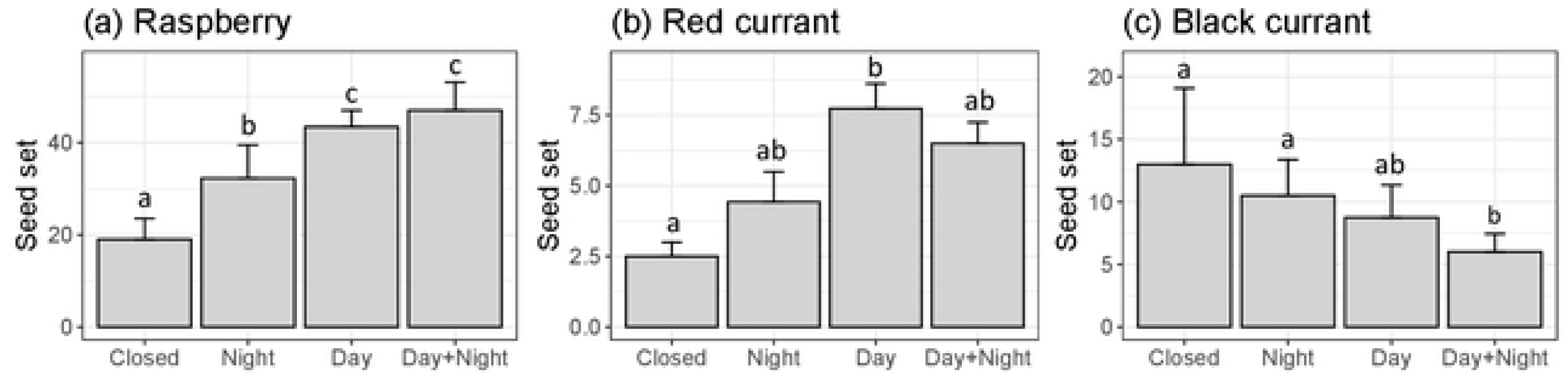
Comparing seed set (mean+se) between Closed, Night, Day and Day+Night pollination treatments for (**a**) raspberry, (**b**) red currant, and (**c**) black currant. Different small letters indicate significant differences with a risk of 5%.

## Discussion

Our results suggest that nocturnal pollinators provide an important contribution to pollination in berry crops, in particular for raspberry, on our study site. Indeed, we report that this crop’s seed set is improved by exposure to nocturnal pollinators, compared to the absence of any pollinator visits, confirming a nocturnal pollination service for this crop. We recognise that our experiment would benefit from a higher sample size and spatial replicates. However, it is noteworthy to specify that such a pollination experiment on day vs. night exclusions is very rare in the literature, due to the challenge in its set up in the field. This experiment requires a presence on site very early in the morning (at sunrise) and also very late in the evening (at sunset). These logistical constraints strongly limit the possibilities of spatial replications and large sample sizes. Moreover, our results follow findings from studies carried out on other crops such as avocado [28], kiwi [52], apple [38], strawberry [39] and lowbush blueberry [53], leading to increasing evidence of nocturnal contributions to crop pollination.

Although a few studies have reported nocturnal pollination services to crops, these remain understudied and their impact on crop production underestimated [13,28]. This is partly due to the fact that nocturnal pollinator monitoring requires field observations at night, which we were not able to do in the present study. Although challenging, the studies that have performed such nocturnal surveys confirmed the presence of nocturnal crop pollinators, such as Lepidoptera, Coleoptera and Diptera [37,52]. To overcome these challenges, new technological methods are being developed to allow night-long surveys to be performed, such as camera traps specifically designed to capture plant-pollinator interactions [54]. Once the nocturnal pollinator community is characterised, single visit pollen deposition techniques, widely used in diurnal pollination studies (see [55] for an example) can be co-opted to evaluate nocturnal pollinator efficiencies. Using this method, a recent study demonstrate that pollen deposition at night varies between pollinator taxa, emphasising the importance of understanding the pollinating efficiency of different nocturnal pollinators [52].

The general pattern we observed for fruit and seed set for all three crops suggests that pollination service is higher for diurnal than for nocturnal visits, and maximised by the combination of diurnal and nocturnal visits. This further highlights that without nocturnal pollinators maximal crop yields cannot be reached. However, the differences between pollination treatments were not all statistically significant. Indeed, we report significant nocturnal pollinator contribution to crop pollination only for raspberry seed set, but not for fruit or seed set of red currant or black currant. The mesh netting used for pollinator exclusion did not prevent wind and self pollination, which could explain why there were few significant effects, in particular for black currant which is predominantly self-fertile [43]. Moreover, for black currant we find a reverse pattern of seed set compared to the other crops. This is due to the intrinsic link between fruit set and seed set. Indeed, black currant fruit set success rate was low, particularly in the Closed pollination treatment, leading to a 5-fold reduction in sample size. Our study also confirms the benefits of the insect-mediated pollination services for crop yield of raspberry and red currant [2], in particular the service provided by day-active pollinators, which increased raspberry fruit set and red currant fruit set and seed set. Interestingly, our pollinator observations showed that while raspberry flowers were visited by an abundant pollinator community, red currant flowers received very few insect visits. Here, our observation time was limited to 5-minute sessions, meaning we may have observed a snippet of all diurnal pollinators responsible for this pollination service, in particular for red currant. Using innovative approaches such as camera traps [54] could improve upon this by recording pollinator visits over periods of 24 hours, and thus allow us to observe all diurnal and nocturnal flower visits.

Our results should however be interpreted with caution since we had few plant replicates. In future research, similar studies with larger sample sizes could allow to take plant identity into account as a random effect, to untangle the effects of treatment and plant individual. This would also increase the floral display and thus increase the number of pollinators visiting flowers. Moreover, challenges in setting up such a day vs. night pollination experiment prevented us from replicating the study on different sites. Future studies could expand upon our observations, by examining nocturnal pollination services at larger scales with more plants and sites along a gradient of landscape complexity (i.e. from semi-natural areas to intensive agricultural landscapes) to explore how less favourable conditions for diverse pollinator communities would affect nocturnal pollination services.

The decline of diurnal pollinators and its causes are well documented [7,56]. However, since knowledge about nocturnal pollinators is so sparce, threats to them remain largely unknown [33]. In particular, Artificial Light At Night (ALAN) is known to affect biodiversity from the individual organism to the ecosystem scale [57]. Regarding pollinators, one study found that light pollution reduced the amount of pollen transported by moths [58]. Similarly, ALAN has been found to disrupts nocturnal pollinator networks, with negative knock-on effects on plant reproduction [59]. Research should focus on understanding the effects of ALAN on crop pollination, particularly in areas where light pollution is intense, like cities, since there is an increasing demand for agricultural land to be close to where people live [60]. A recent study found that urbanisation affects diversity of pollen transported by nocturnal pollinators [36]. Production of crops grown in urban landscapes will likely be deterred by urbanisation, and in particular by ALAN, which disrupt pollinator communities.

In agricultural landscapes, an additional threat to nocturnal pollinators currently is the application of pesticides at night. Indeed, pesticide guidance labels, government institutions, and universities recommend that certain pesticides (especially insecticides) not be applied when bees are active, often recommending application either in the early morning or in the evening [61,62]. In particular, in France, the recent update of the “pollinator-friendly” legislation prevents pesticide applications during the day, but allows it at night [63]. These recommendations stem from a lack of knowledge of the existence and importance of nocturnal pollinators, reinforcing our argument that more research on nocturnal pollinators is needed, to safeguard the pollinators themselves and the valuable pollination services they provide to crops.

In conclusion, our study highlights the contribution of nocturnal pollinators to raspberry crop production, and points to a potential contribution of nocturnal pollinators to yield of black currant and red currant. We suggest that only the combination of both diurnal and nocturnal pollinator visits can maximise crop production. This study is, to our knowledge, the first to evaluate nocturnal pollination services for these crops, adding to a growing body of evidence that nocturnal insects contribute to crop pollination services. Since nocturnal pollination is understudied, threats to night-active pollinators are poorly described compared to their diurnal counterparts. Thus, we highlight the urgency for nocturnal pollinators to be considered in future research and management practices.

## Acknowledgements

This study was carried out during a period of confinement due to the COVID-19 pandemic. We would like to thank Pascal and Doris Requier for accepting this experiment in their orchard, and Noelia Martinez Sartore for help with field work.

## Data accessibility

The data presented in this manuscript are available through the figshare repository https://doi.org/xxxx/m9.figshare.xxxxx [to be complete before final acceptance].

## Author Contribution

**Conceptualization**: Fabrice Requier

**Data Curation**: Fabrice Requier

**Formal Analysis**: Elsa Blareau and Fabrice Requier

**Investigation**: Fabrice Requier

**Methodology**: Fabrice Requier.

**Project administration**: Fabrice Requier

**Resources**: Fabrice Requier

**Supervision**: Fabrice Requier

**Validation**: Fabrice Requier

**Visualization**: Elsa Blareau and Fabrice Requier

**Writing – Original Draft Preparation**: Elsa Blareau and Fabrice Requier.

**Writing – Review & Editing**: Elsa Blareau and Fabrice Requier

